# NMDA receptor in vasopressin 1b neurons is not required for short-term social memory, object memory or aggression

**DOI:** 10.1101/670893

**Authors:** Sarah K. Williams Avram, Jarrett Fastman, Adi Cymerblit-Sabba, Adam Smith, Matthew Vincent, June Song, Heon-Jin Lee, Michael C. Granovetter, Su-Hyun Lee, Nick Cilz, Michelle Stackmann, Rahul Chaturvedi, W. Scott Young

## Abstract

The vasopressin 1b receptor (Avpr1b) plays an important role in social behaviors including aggression, social learning and memory. Genetic removal of Avpr1b from mouse models results in deficits in aggression and short-term social recognition in adults. Avpr1b gene expression is highly enriched in the pyramidal neurons of the hippocampal cornu ammonis 2 (CA2) region. Activity of the hippocampal CA2 has been shown to be required for normal short-term social recognition and aggressive behaviors. Vasopressin acts to enhance synaptic responses of CA2 neurons through a NMDA-receptor dependent mechanism. Genetic removal of the obligatory subunit of the NMDA receptor (Grin1) within distinct hippocampal regions impairs non-social learning and memory. However, the question of a direct role for NMDA receptor activity in Avpr1b neurons to modulate social behavior remains unclear. To answer this question, we first created a novel transgenic mouse line with Cre recombinase knocked into the Avpr1b coding region to genetically target Avpr1b neurons. We confirmed this line has dense Cre expression throughout the dorsal and ventral CA2 regions of the hippocampus, along with scattered expression within the caudate-putamen and olfactory bulb. Conditional removal of the NMDA receptor was achieved by crossing our line to an available floxed Grin1 line. The resulting mice were measured on a battery of social and memory behavioral tests. Surprisingly, we did not observe any differences between Avpr1b-Grin1 knockout mice and their wildtype siblings. We conclude that mice without typical NMDA receptor function in Avpr1b neurons can develop normal aggression as well as short-term social and object memory performance.

**Significance Statement:** Activity of neurons that express vasopressin 1b receptor are essential for aggressive and social recognition behaviors. We created a novel transgenic mouse to allow selective targeting of vasopressin 1b neurons. Our studies indicate that NMDA receptor expression in vasopressin 1b neurons (including most CA2 neurons) are not required for development of the typical expression of aggression or recognition memory. Thus, CA2 neurons may have a unique way of incorporating novel stimuli into memory that deserves further investigation.

## Introduction

The neuromodulatory actions of arginine-vasopressin (Avp) are required for the typical expression of social and stress-related behaviors in clinical populations and preclinical models (Caldwell, Aulino et al. 2017, Williams Avram and Cymerblit-Sabba 2017). Avp acts in the central nervous system through its G-protein coupled receptors, Avpr1a and Avpr1b. Recent data indicate that variation in Avpr1b signaling may have maladaptive impacts on human social behavior. Avpr1b SNP variants are positively correlated with autism diagnoses (Francis, Kim et al. 2016) and with increased emotional aggression (Luppino, Moul et al. 2014). Similarly, life-long disruption of Avpr1b signaling through genetic removal in knockout (KO) mice results in altered social aggression and social memory performance (Wersinger, Ginns et al. 2002), while leaving spatial and object memory performance and olfactory discrimination intact, indicating a special role in the social aspect of memory (Wersinger, Ginns et al. 2002, Wersinger, Kelliher et al. 2004, Wersinger, Caldwell et al. 2007, Caldwell, Wersinger et al. 2008). Aggression in Avpr1b knockout mice was decreased without co-occurring deficits in predatory aggression or anxiety and remained deficient in a KO on a more outbred strain of mice (Wersinger, Caldwell et al. 2007, Wersinger, Temple et al. 2008, Caldwell and Young 2009, Caldwell, Dike et al. 2010). Furthermore, decreases in social aggression occurred in both sexes, suggesting a conserved signaling mechanism for males and females, unlike many of Avp’s actions through Avpr1a (Terranova, Song et al. 2016, Williams Avram and Cymerblit-Sabba 2017).

Avpr1b shows a distinct anatomical expression profile from Avpr1a. Mapping of Avpr1b mRNA expression in mice revealed the highest levels in the pyramidal neurons of the dorsal CA2 hippocampal region, with a few cells expressing in the medial amygdala and paraventricular nucleus of the hypothalamus (Young, Li et al. 2006). Pagani and colleagues used lentiviral delivery of the Avpr1b gene into the dorsal CA2 of Avpr1b KO mice and were able to rescue the aggressive phenotype, indicating Avpr1b expression in the dorsal CA2 is essential to expression of species-typical aggressive behavior (Pagani, Zhao et al. 2015). Furthermore, optogenetic activation of vasopressinergic neuron terminals in the dorsal CA2 enhances and extends social memories (Smith, Williams Avram et al. 2016). Local pharmacological antagonism of Avpr1b prevents this extended social memory, suggesting Avpr1b activity in the dorsal CA2 is required for the effect. However, the molecular mechanism through which Avpr1b may be influencing neuronal plasticity to contribute to these behavioral functions is unknown.

Avp application results in extended increased EPSC amplitude in dorsal CA2 neurons, a response that is blocked by NMDA glutamate receptor antagonists (Pagani, Zhao et al. 2015). Furthermore, the increased EPSC amplitude was not mediated by altered local inhibitory signals as bicuculline had no effect on this response. This indicates that NMDA receptors may be critical for Avp’s role in modulating dorsal CA2 neural activity. NMDA receptor signaling is required for plasticity in postsynaptic response in many cell types (Baez, Cercato et al. 2018). The NMDA receptor is a tetrameric ion channel composed of two obligatory glutamate receptor ionotropic NMDA type 1 (Grin1) subunits and 2 subunits from the Grin2 or Grin3 families, the composition of which varies with cell-type and throughout development (Hansen, Yi et al. 2018). Genetic removal of Grin1 gene results in deficits in synaptic plasticity and learning (Zweifel, Argilli et al. 2008, Hansen, Yi et al. 2018).

To determine the role that plasticity may have in Avpr1b neuron-dependent behaviors, we selectively inactivated NMDA receptors in Avpr1b neurons through genetic removal of the obligatory Grin1 gene, and assessed aggression, memory and anxiety-like behaviors. We predicted that these KO mice (Avpr1b^Grin1-/-^) would exhibit impairments in social memory performance and aggressive behaviors. Surprisingly, we did not observe any behavioral impact in any measure in the Avrp1b^Grin1^ mice, suggesting that a chronic loss of the Grin1 subunit in Avpr1b neurons is well tolerated with regard to social aggression and memory.

## METHODS

### Generation of Avpr1b-Cre Knock-in Mice

To generate the targeting vector construct (Figure 1A), a BAC plasmid clone (AC120217) that contains the mouse Avpr1b genomic locus was used to amplify various genomic segments by PCR. A 4 kb 5’ fragment containing the Avpr1b promoter and part of exon 1 with its start codon was inserted into a targeting vector (pGKneoF2L2 DTA, kindly donated by Dr. Philippe Soriano at Fred Hutchinson Cancer Research Center, MIT, Boston, MA) upstream of a phosphoglycerate kinase promoter-driven neomycin resistance cassette (pgk-neo^r^ or neo) flanked by FRT sites. The Cre-NLS (from the pBS185 plasmid from the former Life Technologies, Gaithersburg, MD) and SV40 poly A signal (from the pTRE-tight plasmid from Clontech Laboratories, Mountain View, CA) sequences were cloned right after the 5’ homology fragment. For the 3’ homology part of the construct, a 3.2 kb of fragment containing intron 1 was inserted between the neo and pgk-DTA cassettes (diphtheria toxin A gene), which were used to select against random insertion. Following electroporation into ES cells (LC3 cell line; (Tra, Gong et al. 2011)), DNA from G418 resistant clones was screened by PCR and digested with XbaI for analysis by Southern blotting using an internal probe. Correctly targeted alleles produced a 7.2 kb XbaI fragment owing to the insertion of the selectable drug-resistant maker (neo gene), compared to the wildtype (WT) allele of 9.8 kb (Figure 1B). The positive ES cells were injected into blastocysts to generate chimeras. Avpr1b-Cre heterozygous mice (Hets) were then crossed with a transgenic mouse expressing FLPe recombinase using a human beta-actin promoter (B6;SJLTg(ACTFLPe)9205Dym/J; The Jackson Laboratory, Bar Harbor, ME) to remove the neo cassette that was flanked by FRT sites (Rodríguez, Buchholz et al. 2000). These Avpr1b-Cre heterozygotes (Avpr1b-Cre^+/-^) were then either bred together or with C57Bl/6J partners. Mice used in gene expression these experiments were from litters backcrossed at least 3-4 generations into the C57Bl/6J strain.

**Figure 1.**
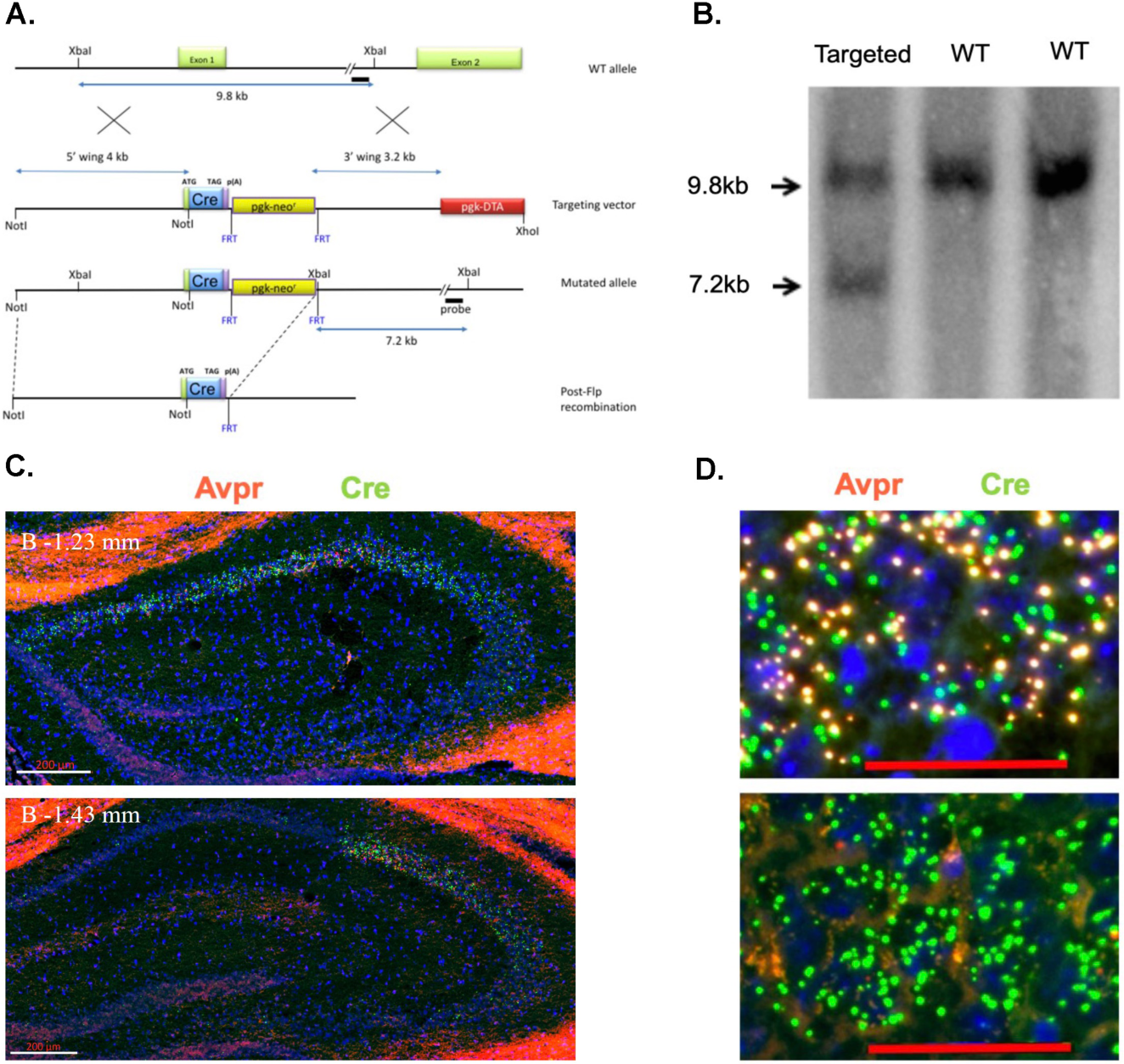
A. An illustration of the design of the targeting vector construct indicating the portion of the Avpr1b gene surrounding exon 1 that was replaced through homologous recombination. B. Southern blot indicating the Cre recombinase gene was knocked in to the appropriate locus (genomic DNA restricted with XbaI). C. Representative images of double in situ hybridization histochemistry on sections (16μm) from a heterozygous mouse demonstrating Cre recombinase mRNA is selectively expressed in neurons that co-express Avpr1b mRNA along the rostral-caudal axis (n=3 mice). Avpr1b transcripts locations are indicated by orangish-red dots and Cre by green dots. (bar=200μm) D. Representative magnified images of (top) co-expression of Cre-recombinase and Avpr1b mRNA in the heterozygous mouse and (bottom) lack of Avpr1b transcripts in the homozygous mouse (bar=50μm).

Cre-dependent TdTomato reporter mice (Ai9) were purchased from Jackson Laboratory (Strain Name: B6.Cg-Gt(ROSA)26Sortm9(CAG-tdTomato)Hze/J, Stock Number: 007909). These mice were bred to Avpr1b-Cre^+/-^ mice, resulting in Avpr1b-Cre^+/-^;TdTomato double transgenic offspring. Two adult males and females were used to assess Avpr1b-Cre distribution.

Mice containing loxP sites flanking the Grin1 (Grin1^lox/+^) gene were purchased from Jackson Laboratory (Strain Name: B6.129S4-Grin1^tm2Stl^/J, Stock Number: 005246) (Tsien, Chen et al. 1996) and backcrossed 2-3 generations into the C57Bl/6J strain prior to breeding. Following this, heterozygotes were then bred together to maintain the colony. To generate experimental animals, male mice with a confirmed genotype of fully floxed Grin 1 and wildtype Avpr1b (Grin1^lox/lox^: (Avpr1b^+/+^) were mated to females that were genotyped heterozygous floxed Grin1 and heterozygous for the Avpr1b-Cre (Grin1^lox/+^:Avpr1b^+/Cre^). The resulting offspring had the following genotypes 1) Grin1^lox/lox^:Avpr1b^+/+^, 2) Grin1^lox/+^:Avpr1b^+/+^, 3) Grin1^lox/+^:Avpr1b^+/Cre^ 4) Grin1^lox/lox^:Avpr1b^+/Cre^. The first two genotypes were labeled as wildtype (WT). The third, heterozygous group were labeled as Avpr1b^Grin1+/-^. The last group was labeled as the Grin1 knockout Avpr1b^Grin1-/-^. Animals for behavioral experiments came from a total on 6 different breeding pairs.

Stimulus mice used in social recognition testing were sexually naïve, ovariectomized Balb/c adult females. They were purchased from Jackson Labs (Stock # 000651) and ovariectomied. Briefly, a small dorsal midline incision was made, the muscle wall spread using forceps, and the ovaries were removed. Following a two-week recovery period, females were singly housed for at least one week prior to testing. Stimulus mice used in the social habituation and social novelty tasks were adult sexually-naïve group-housed C57BL/6J males bred in our colony. On an annual basis, new breeders are acquired from Jackson Labs (Stock # 000664). Stimulus mice for the aggression testing were adult sexually-naïve group-housed Balb/c males (Stock # 000651). Stimulus mouse choice for each test is based on extensive pilot work in our lab showing the strongest and most consistent response from group and single-housed male C57Bl/6J mice. Stimulus mice were used only once per day.

### Genotyping

Mice were genotyped by PCR using DNA extracted from tail snips. A single forward primer, V1BR#9 (GAAACGGCTACTCTCTCCGATTCCAAAAGAAAG), was designed for amplification of both WT and recombined loci. The first reverse primer, V1BR#5 (ACCTGTAGATATTTGACAGCCCGG), was designed to amplify the WT loci (762 bp PCR product). The second reverse primer, Cre.c35 (GATATAGAAGATAATCGCGAACATCTTCAGGTTCT), was designed to detect the Cre recombinase transgene (679 bp). PCR was carried out for 40 cycles with denaturation at 94°C, annealing at 60°C, and extension at 72°C, all for 1 min. Grin1 mice were genotyped using a forward primer NR1loxP2 (ACAATAGAGATTCAAGGCTGATCAAGG) and reverse primer NR1loxP3 (CTCTGGGTGGCTTGCCTGGCTGTATGTT). PCR was carried out for 40 cycles with denaturation at 94°C, annealing at 60°C, and extension at 72°C, all for 45 s. TdTomato mice were genotyped for the wildtype band using a forward primer, oIMR9020, (AAGGGAGCTGCAGTGGAGTA) and reverse primer, oIMR9021, (CCGAAAATCTGTGGGAAGTC). The mutant band is genotyped using the forward primer, oIMR9105, (CTGTTCCTGTACGGCATGG) and reverse primer, oIMR9103(GGCATTAAAGCAGCGTATCC). PCR was carried out for 40 cycles with denaturation at 95°C, annealing at 65°C, and extension at 72°C, all for 45 s

### RNA Chromogenic *In Situ* Hybridization Histochemistry

For in situ hybridization histochemistry studies, mice were anesthetized with isoflurane, decapitated and the brains removed and frozen on powdered dry ice within 5 minutes. Brains were sliced on a cryostat and 16um sections were collected onto slides and kept frozen until the time of assay. Avpr1b and Cre recombinase transcripts were simultaneously detected in fresh-frozen brain sections from heterozygous Avpr1b-Cre mice using the Affymetrix ViewRNA duplex kit (Catalog number: QVT0013; Santa Clara, CA) as previously described (Young, Song et al. 2016). Alternate slices were used to detect Grin1 and GAD expression. The probe sets targeted Avpr1b (catalog number: VB1-16867), Cre recombinase (VF6-16306), Grin1 (VB1-14161) and GAD (VB6-12632).

### Collection of Avpr1b-tdTomato Brains

Adult male and female Avpr1b-Cre^+/-^;TdTomato animals were transcardially perfused with 4% paraformaldehyde. Brains were removed and postfixed for 24 hours. Following an overnight rinse in 1M phosphate buffer saline solution, brains were transferred to a 30% sucrose solution. Brains were sliced at 50um on a cryostat (Leica3050) into cryoprotectant and mounted onto charged slides. Slides were stained with DAPI (300nM) and coverslipped with polyvinyl alcohol/1,4 diazabicyclo[2.2.2]octane/glycerol solution (PVA-DABCO).

### Imaging and Image Analysis

All sections were imaged using the Zeiss AxioScan Z1 slide scanner and online stitching and shading correction using a 20X objective. Image contrast was adjusted for presentation in Zen Blue. Image analysis was performed in Arivis Vision 4D software. For Avpr1b gene expression in the Avpr1b-Cre line, regions of interest were drawn over the dorsal CA2 and fasciola cinereum (FC). mRNA puncta were segmented based on intensity threshold and shape. Puncta were counted and presented as number per area. Measurements were made bilaterally on 3 sections/mouse. Measurements were then averaged per region, and then averaged per mouse. For Grin1 gene expression, mean intensity of Grin1 label was quantified. Regions of interest were drawn using the nuclear counterstain (DAPI) channel. Regions were drawn around areas of strong Avpr1b expression including the CA2 and fasciola cinereum (FC). Adjacent CA1, CA3, dentate gyrus and layer 2 motor cortex were also measured. Measurements were made bilaterally on 3 sections/mouse. Measurements were then averaged per region, normalized to cortical measurements and averaged per animal. Three animals in each genotype measured.

### Mouse Housing Conditions

All housing and procedures were approved by the Animal Care and Use Committee of the National Institute of Mental Health. Mice were housed in an AAALAC accredited specific pathogen-free vivarium kept at constant temperature and humidity (∼21°C, 50%), in plastic micro-isolator cages (12”×6.5” ×5.5”) containing wood chip bedding (Nepco Beta-chips) and cotton nestlets. All cages were maintained on high-density ventilated racks (Super Mouse 750, Lab Products Inc.). Mice were maintained on a 12-h light cycle (lights off at 1500h) with *ad libitum* access to standard mouse chow (Purina Lab Diet, Product #5R31) and water bottles. Cages were changed on a bi-weekly basis primarily by the same animal caretaker. All breeding pairs were fed a high-fat diet (Purina Lab Diet, Product #5058) as a measure to reduce pregnancy and pup loss. All offspring were weaned at ∼21 days of age into cages with their same-sex littermates. No animals that were singly housed during adolescence were used in behavioral testing, but may have been used for gene expression studies. All animals used in behavioral experiments were adult males that remained group-housed with littermates for the entirety of testing. Testing began when the animals were 90-120 days old.

### Behavioral Procedures

#### Experimental Design

All mice went through the same series of behavioral tests in the following order: Elevated O-Maze, Social Recognition Novel-Familiar, Social Recognition Novel-Novel, Social Habitation-Dishabituation, Object Habituation-Dishabituation, Olfactory Habituation-Dishabituation, Social Novelty Preference, and following a 2-week period 3 tests of aggression. For the elevated O-maze test, each mouse was measured sequentially, while the recognition, habituation, and aggression tasks were parallelized to allow four mice to be tested simultaneously. For simultaneous testing scenarios, all test spaces were assessed for even illumination. All behavioral tests were separated by 48-72 hours. Three small groups of animals were run through the battery of tests. The groups were separated by several months, however, each group contained representatives from all genotypes. The total number of animals in each group was WT= 9, Avpr1b^Grin1+/-^ = 8, Avpr1b^Grin1-/-^ = 11. All testing was performed during the light phase of the cycle. All testing was performed in a room separated from the colony room.

#### Elevated O-Maze

Anxiety-like behavior was tested in an elevated O-maze (San Diego Instruments, San Diego, CA). Testing took place in a darkened room (20 lux). A row of LED lights controlled by a dimmer switch was set to illuminate only the open arms at an intensity of 60 lux at the maze surface. All mice were moved into an anteroom of the behavioral suite 30 min prior to testing. Mice were placed on the open arm of the maze facing a closed arm and were allowed to explore the maze for 5 min. Mice were recorded using a ceiling mounted camera connected to a Dell computer running Ethovision software (Noldus Information Technology, Leesburg, VA). Trials began as soon as the experimenter left the room.

#### Social Recognition Testing

For the two-trial recognition test, mice were brought to the testing room and habituated to a fresh mouse cage containing clean bedding for 30 min. An unfamiliar ovariectomized Balb/C female was placed in the cage and allowed to freely interact with the experimental male for 5 min. Then the female was returned to her home cage. The experimental animal remained in the testing cage. Following a 30-min interval, the experimental mouse was re-exposed to either the original or a novel female for another 5-min interval.

#### Social Habituation-Dishabituation

Mice were brought to the testing room and habituated to fresh cages for 30 min. A group-housed adult C57Bl/6J stimulus male was added to the cage for four 1-min trials of unrestricted interaction with an inter-trial interval of 3 min. During the intervals, the stimulus mouse was kept in a holding container on an adjacent table and the experimental animal remained in the testing cage. The fifth trial introduced a novel stimulus male from a different home cage.

#### Object Habituation-Dishabituation

This procedure is similar to the social habituation protocol, except novel objects were placed in the cage. Objects were placed into the same corner of the cage for each trial. Location of the corner was randomized. The objects were small brightly colored plastic items with complex shapes (small water bottles, rubber duckies, scintillation vials with colored fluid). All items were cleaned with ethanol and water prior to use. All items were previously found to elicit similar amounts of initial investigation.

#### Olfactory Habituation-Dishabituation

Olfaction was tested using a habituation-dishabituation task adapted from one we have used previously (Lee, Caldwell, Macbeth, Tolu, et al. 2008b). Mice were placed in a clean cage containing fresh bedding. Following a 30-min habituation, three odorants were presented for 1 min, three times each, with a 3-min inter-trial interval: water, almond (1:100), male mouse urine (1:100), for a total of nine trials. Presentation was made with 100 ul of odorant solution added to cotton balls placed in metal strainer spheres. The amount of time mice spent with their snout in proximity (1 cm or less) to the metal strainer was recorded. Odorant solutions were diluted in distilled water from stock solutions. Almond scents were extracts (McCormick and Company Inc, Spark MD) and urine samples were mixtures from several adult male mice. Urine was collected by placing animals in modified mouse metabolic chambers that have a grid mesh floor with a funnel to collect liquid below. Individual animals were allowed to stay in the chamber for 30 minutes. Urine samples were combined, flash frozen on dry ice and stored at -80°C. Urine was thawed, diluted, and kept at 4°C between testing days.

#### Social Novelty Preference

Test mice were brought to the experimental room and allowed to acclimate for 30 min. Test mice were placed in the center of the Plexiglas three-chambered apparatus (each chamber 18×45×30cm) for a 5-min habituation phase, followed by a 5-min choice phase. The side chambers contained either a male littermate who had been group-housed with the experimental mouse or a novel age- and weight-matched C57BL/6J male mouse. Stimulus mice were wearing collars and leashes that were affixed to the corner of the chambers. This prevented the stimulus mouse from leaving its designated chamber but allowed for full access to the stimulus mouse by the test animal (adapted from (Winslow 2003). The collars were small beaded plastic zip ties affixed to small four-inch chains mounted on the wall of the chamber. The chains were mounted to the wall most distal to the entry to the chamber. The length of the tether allowed the stimulus mouse to access half of the chamber. Stimulus mice were habituated to the collars for 15-30 minutes prior to being placed in the chamber.

#### Aggression

Following two weeks of isolated housing, subjects were tested on 3 occasions, always 48-72 hours between testing sessions. Subjects were brought to the testing room (∼ 70 lux, with 60dB white noise), between 9:00 AM-1:00 PM, weighed, and returned to their home cage without food or water for 30 minutes. Adult, weight-matched sexually-naïve, group-housed Balb/C male mice were used as intruders. Intruders were placed on the opposite side of the cage. Experimental males were given 5 min to exhibit aggressive behavior. If aggressive behavior was observed, the latency was noted, and the intruder removed after a 2 minute period. Stimulus males were never used more than once on the same day and never with the same subject on a separate testing session. Intruder males were no longer used after they had been attacked five times. If an intruder male was observed initiating an attack on a subject, he was removed from the test and no longer used as an intruder. Videos were recorded with a Panasonic HDC-TM700 camera and scanned by an observer blind to the identity of the mouse for the initial instance of aggression defined as the first bite observed from the subject coded.

#### Behavioral Analysis

Videos of behavioral tests were recorded from above and coded by an observer blind to the identity of the mouse using JWatcher Software (http://www.jwatcher.ucla.edu/ (Blumstein, Daniel et al. 2010)).

#### Behaviors analyzed for social behavior (aggression, social recognition, social habituation-dishabituation) tests included

<u>1)</u> *Sniff anogenital:* the mouse sniffs the anogenital region of the stimulus, including the base of the tail. The mouse may approach from behind the stimulus mouse or burrow its head under the abdomen of the stimulus to place its nose between the hindlimbs. Sniffing may occur while mice are locomoting, where the mouse follows closely behind the stimulus, keeping its nose in contact with the anogenital region of the stimulus. 2) *Aggression:* the mouse attacks the stimulus mouse by lunging forward, biting, and often thrashing. When two mice are fighting, the mouse that initiated an attack is considered the aggressor. Attacks are commonly foreshadowed by circling (aggressor mouse runs circles around victim with head oriented toward victim) and/or tail rattling (aggressor mouse raises and rapidly shakes its tail while facing toward victim). 3) *Allogrooming:* the mouse licks the fur of another mouse (often rapidly) or uses its forepaws to comb the fur of another mouse. Grooming is usually focused on the back, shoulders and head of the recipient mouse, and the recipient mouse may either continue to behave freely or freeze and remain motionless while being groomed. 4) *Mounting:* the mouse approaches another mouse from the rear and places its forepaws on the back of the recipient mouse while thrusting its lower body forward. The recipient mouse will often attempt to flee, with the test mouse following closely behind and attempting to re-mount. General nonsocial behaviors were also measured. 5) *Autogrooming:* while sitting still, the mouse licks its fur, swipes its face with its forepaws, and/or uses any limb (though usually hindlimbs) to scratch itself. 6) *Digging/burrowing:* the mouse uses its face or limbs to actively displace bedding. The mouse may bury its face in bedding while sitting still or push bedding in front of it while moving using its head and forepaws. 7) *Climbing:* the mouse moves to any surface other than the floor of the cage or testing chamber. Climbing may occur on the lip of the cage or testing chamber, on the metal air vent attached to the cage, or on top of a corral or other object within the cage. The mouse may be still or in motion while climbing. 8) *Rearing:* the mouse raises its forepaws off of the floor and sits or stands on its hindlimbs. The mouse may sit on its haunches with its forepaws in the air, or stretch with its hindlimbs extended and its forepaws resting on an object or on the side of the cage or testing chamber

#### Behaviors analyzed for object and olfactory habituation experiments included

*Sniff:* the mouse sniffs the object, indicated by head movement and snout within 1 cm of object. Sniffing may occur while mice are locomoting. *Climbing, Autogrooming, Digging/burrowing, and Rearing* are scored as described above.

#### Behaviors analyzed for the elevated O-maze included: 1)

*Time in light/dark arms:* the total amount of time spent in each portion of the maze. The mouse must have all four paws within the boundary. 2) *Stretch into Arm:* while in the open arm, the mouse leans forward and stretches its forepaws into the opposite arm. The hindlimbs must remain in the original arm.

#### Statistical analysis

All data was checked for normality using the Shapiro-Wilks test. Avpr1b gene expression was assessed with a student’s t-test. Grin1 mRNA expression was assessed with two-way ANOVA and planned posthoc comparisons. Litter number comparisons used the Kolmogorov-Smirnov test. The elevated O-maze measures were analyzed using one-way ANOVAs and Welch’s test (for non-normal) to compare genotypes. All memory and habituation behavioral measures were analyzed using two-way repeated measures ANOVAs comparing Trial and Genotype, with planned comparisons of genotype in Trial One. Additionally, ratio scores were computed and compared with one-way ANOVAs or Krusal-Wallis (for non-normal sets). Aggression likelihood test used a Chi-square analysis to compare genotypes. Aggression duration and frequency were compared with two-way repeated measures ANOVAs with Geissner-Greenhouse correction for unequal variance. All data analysis was performed in Prism Software (version 8).

## Results

### Validation of Transgenic Mouse Model

#### Cre expression matches Avpr1b expression

Cre recombinase expression was confined to the same cells as Avpr1b gene expression as shown by *in situ* hybridization histochemistry. Avpr1b mRNA and Cre mRNA were seen in dorsal CA2 and immediately adjacent CA3 areas of the hippocampus (Figure 1C), consistent with our previous Avpr1b localization (Young, Li et al. 2006).

#### Avpr1b-expressing neuron localization

To further characterize the extent of Avpr1b gene expression, the Avpr1b-Cre line was crossed with a fluorescent reporter line (male and female, n=2). We see cellular expression in the expected dorsal CA2 and FC neurons (Figure 2A-D), where the majority of the pyramidal cells are labeled. Additionally, we observe labeled neurons in ventral hippocampal areas (Figure 2E).

**Figure 2.**
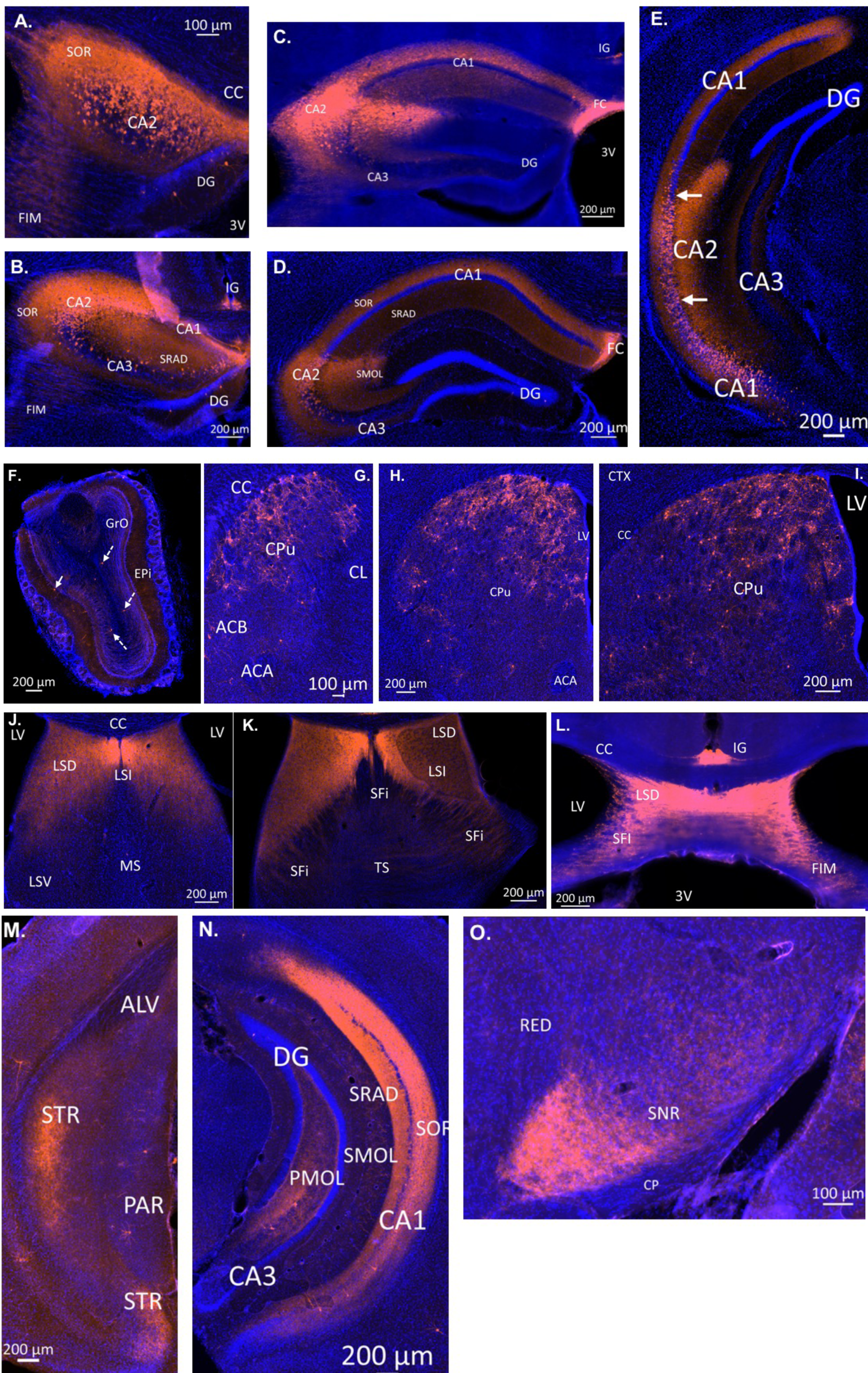
Avpr1b-Cre expression throughout the adult brain. A-I) Location of neuronal cell bodies expressing Cre-dependent TdTomato. Cells are observed throughout the anterior-posterior CA2 (A-E) and FC (A-D). Scattered cells are observed in the olfactory bulb (F). Cells are observed throughout the anterior-posterior sections of the caudate putamen (G-I). Indusium Griseum neurons can be observed quite anterior (L) and continued posterior until the middle of the hippocampus (C). Neuronal projections can be observed within the hippocampus. Dense fibers are observed in the stratum oriens and stratum radiatum in every level where cell bodies exist (A-E). Dense fiber projections continue posteriorly throughout the posterior CA1 (N) and into the transition zone of the subiculum (M). Dense dendritic projections can be observed with the stratum molecular of the CA2 in all layers where cells bodies exist (C-E). In posterior sections dense fibers can be observed within the polymorphic layer of the dentate gyrus (N). Extra hippocampal fibers are observed throughout the lateral septum. The densest projections are found in the LSD and LSI although projections are also observed with in the LSV (J-K). Fibers are also seen in the septofimbral nucleus and throughout the fimbria (L). Dense fibers projections are also observed within the ventral substrantia nigra. CA: cornu ammonis, DG: dentate gyrus, FIM: fimbria, SOR: stratum oriens, CC: corpus callosum, 3V: third ventricle, FC: fasiola cinereum, IG: induseum griseum, SRAD: stratum radiatum, SMOL: stratum molecular, GrO granular olfactory layer: Epi: external plexiform layer, CPu: caudate putamen, ACB: nucleus accumbens, ACA: anterior commissure, CL: claustrum, LV: lateral ventricle, CTX: cortex, LSD: lateral septum dorsal division, LSI: lateral septum intermediate, LSV: lateral septum ventral division, MS: medial septum, SFI: septofimbrial nucleus, TS: triangular septal nucleus:, ALV: alveus of the hippocampus, STR: transition zone of subiculum, PAR: parasubiculum, PMOL: polymorphic layer of dentate gyrus, SNR: substantia nigra reticulata, CP: cerebral peduncle, RED: red nucleus. Blue cells: DAPI, Orange: Cre-dependent TdTomato.

Notably, we observe many labeled neurons in areas that are typically described as ventral CA1. A scattering of neurons is observed within the granule layer of dorsal and ventral dentate gyrus. Neurons of the induseum griseum appear above the corpus callosum for an extended range (∼1 mm, Figure 2B,C,L). Furthermore, we consistently observe numerous scattered cells throughout the caudate-putamen (Figure 2G-I). A few scattered cells are observed within the olfactory bulbs within the granular cell layer and the external plexiform layer (Figure 2F).

Intrahippocampal projections are observed within the dorsal and ventral hippocampus. Specifically, in the dorsal CA1, CA2 and CA3, fibers are most dense in the stratum oriens, with strong projections within the stratum radiatum (Figure 2B-E, N). Dendritic projections are observed in the molecular layer of CA2 and CA1 (Figure 2C-E). Projections within the molecular layer of CA1 continue past the location of cell bodies into the ventral hippocampus and persists to the most posterior region. Moderate projection density is observed in the polymorphic layer of the dentate gyrus in dorsal and denser projections observed more ventrally (Figure 2N). Projections continue posteriorly into the transition zone of the subiculum (Figure 2M)

Extrahippocampal projections are limited to the septum and substantia nigra. Dense fiber projections are observed in the dorsal lateral septum and extending into the septohippocampal nucleus and intermediate lateral septum (Figure 2J-L). Less dense projections are observed in the ventral division of the lateral septum and the triangular nucleus of the septum. Projections are also observed throughout the substantia nigra, with the densest projections observed within the ventral part (Figure 2O).

#### Avpr1b Gene Expression Developmental Timeline

To determine when the Cre recombinase became active during development, we assessed Avpr1b gene expression at postnatal day 1 and postnatal day 7. We observed extremely limited expression within the hippocampus at postnatal day 1 (Figure 3A-B). However, by postnatal day 7 there is abundant expression within the dorsal and ventral hippocampus (Figure 3C-D).

**Figure 3.**
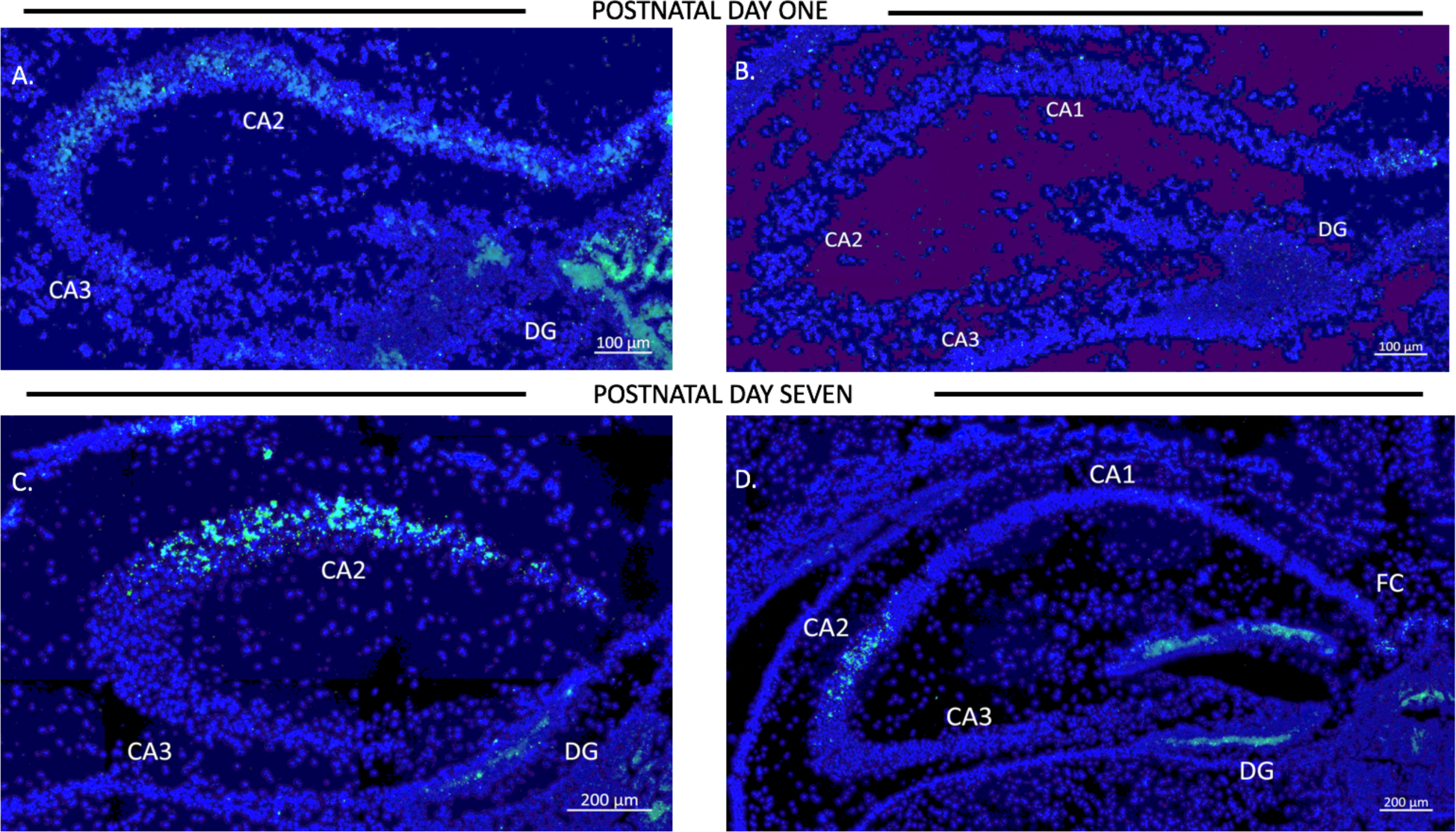
Avpr1b expression throughout development. Representative images of in situ hybridization histochemistry on sections (16μm) from a heterozygous mouse (n=2). A-B. Dorsal hippocampus sections from postnatal day 1 mouse brain along the rostrocaudal axis. Very sparse labeling is observed. C-D. Dorsal hippocampus sections form postnatal day 7 mouse brain along the rostrocaudal axis. Robust Avpr1b mRNA expression is seen across the CA2 and FC regions. Blue: DAPI, Green: Avpr1b.

#### Avpr1b Gene Expression in Avpr1b-Cre animals

To determine if Avpr1b mRNA expression is impacted by the Cre knockin in the dorsal hippocampus, Avpr1b gene expression was measured in the CA2 and FC in WTs (n=4) and Avpr1b-Cre knockins (n=5) (Figure 4A-B). Two-Way ANOVA (Genotype × Region) revealed a significant effect of region [F (1,7)- 44, p< 0.0001], but no effect of genotype [F(1,7)=0.9, p = 0.4].

**Figure 4.**
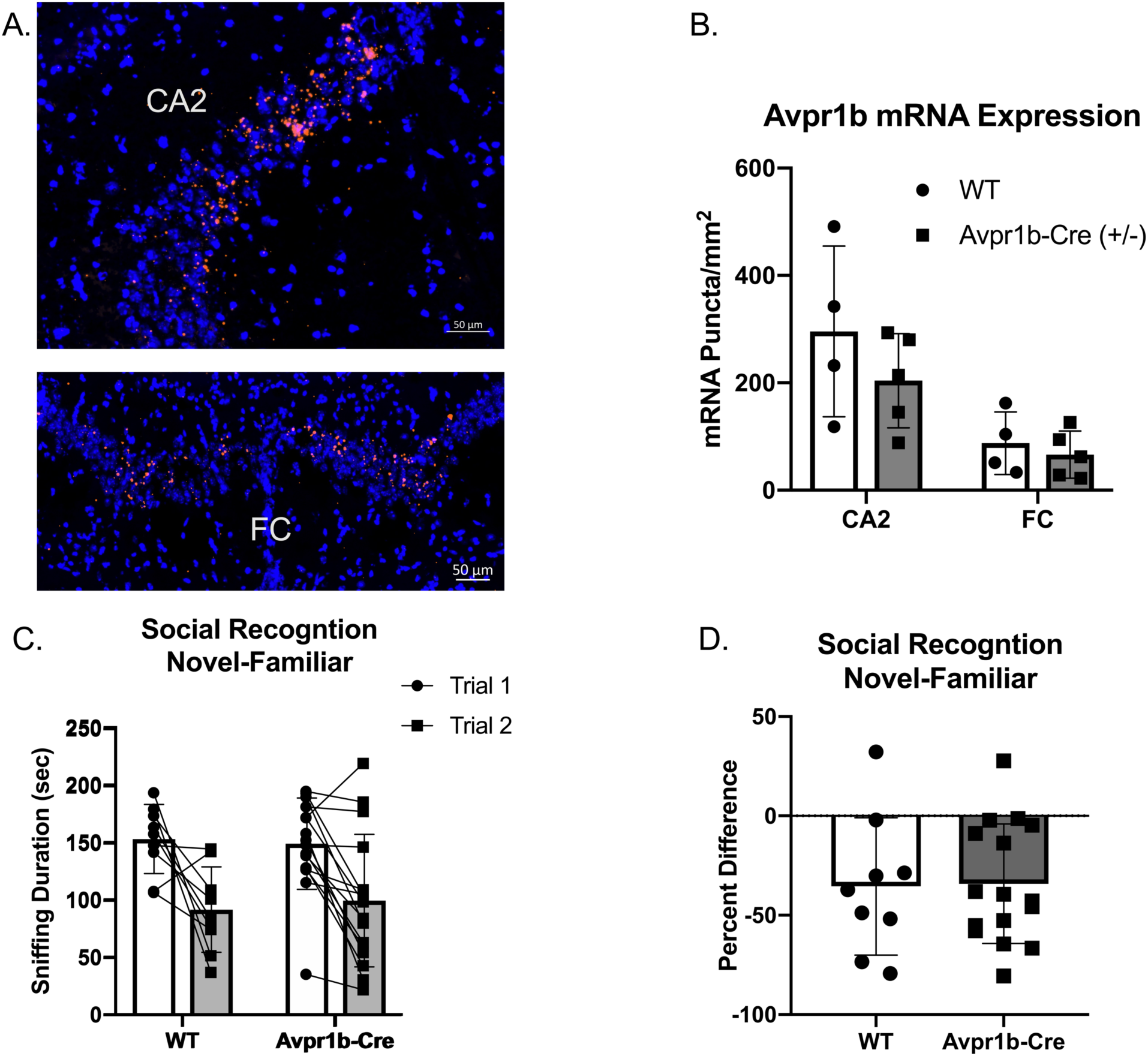
Cre Insertion Does Not Alter Avpr1b Expression or Social Behavior. A. Representative images of in situ hybridization histochemistry of Avpr1b in the CA2 and FC in WT animals. B. Quantification of number of puncta/area. Two-Way ANOVA (Genotype X Region) revealed a significant effect of region [F (1,7)= 44, p< 0.0001], but no effect of genotype [F(1,7)=0.9, p = 0.4]. C. Social interaction and Social Memory are intact. A two-way repeated measures ANOVA revealed a main effect of Trial (F(1,23)= 25.6, p<0.0001), but no effect of genotype. ** indicates difference between Trial 1 and Trial 2 (p< 0.005). D. Ratio of Trial 2 to Trial 1 sniffing. All data presented as individual data points overlaid on means ± SD.

#### Normal growth and Development in Avpr1b-Cre animals

The transgenic knockin did not result in any abnormal breeding occurrences. Specifically, the genotypic ratio was not different than expected [expected; WT =50%, Avpr1b-Cre = 50%; Born: WT= 45%, Avpr1b-Cre = 55%, χ^2^, df =0.50,1,p= 0.5). Litters size was not significantly different than C57’s in our colony [Mean=6.7 ± 2.1 vs 6.03 ±2.4, Kolmogorov-Smirnov=0.18, p= 0.1)]. Furthermore, the adult weights of Avpr1b-Cre males did not significantly differ from WT [Avpr1b: 32.3 ± 3.3, WT: 29.9 ± 2.2; t,df=1.9,22, p = 0.06].

#### Avpr1b-Cre Knock-in does not alter behavior

To assess whether the integration of Cre-recombinase into the Avpr1b locus resulted in behavioral changes, adult male Avpr1b-Cre heterozygous animals were compared with wildtype littermates on the social recognition test (Figure 4C-D). A two-way repeated measures ANOVA revealed a main effect of Trial [F(1,23)= 25.6, p<0.0001], but no effect of genotype [F(1,23)= 0.01, p=0.9]. A planned post hoc comparison of Trial 1 sniffing showed no significant difference in the amount of initial investigation (Sidak’s multiple comparison adjusted p=0.9). The ratio scores also did not differ between the genotypes [t(2) =0.10, p= 0.9].

#### Validation of the Avpr1b^Grin1+/-^ transgenic line

Grin1 mRNA was quantified throughout the dorsal hippocampus where the expression of Cre was most prominent and localized (Figure 5). A two-way ANOVA comparing region and genotype revealed a main effect of region [F(4,35) =7.0, p=0.003], and a main effect of genotype [F(2,35)= 11.1, p=0.0002]. Post-hoc Tukey’s tests revealed that Avpr1b^Grin1-/-^ mice are significantly different from Avpr1b^Grin1+/-^ (p=0.005) and WT (p=0.008) groups in the CA2. Avpr1b^Grin1-/-^ mice are significantly different from Avpr1b^Grin1+/-^ (p=0.007) and WT (p=0.03) groups in the FC. No differences were observed in the CA1, CA3 or dentate gyrus. We confirmed the removal of Grin1 from only pyramidal cells within CA2 and FC regions, as Grin1 was still colocalized with GAD mRNA expression (Figure 5E-F).

**Figure 5.**
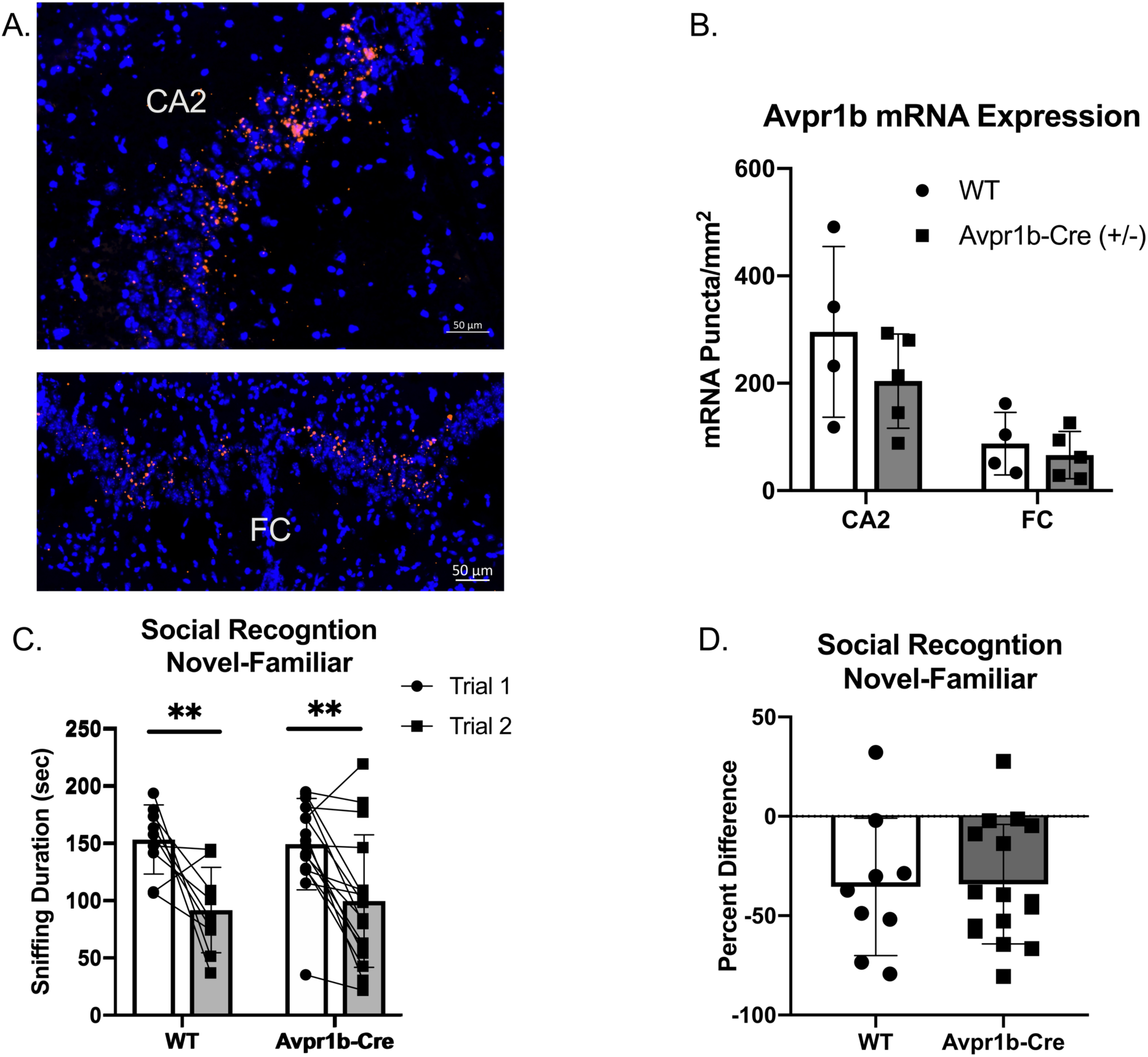
Grin1 is Reduced in Transgenic Mice. Double in situ hybridization of Avpr1b (green) and Grin1 (red) mRNA in the dorsal hippocampus in WT (A) and Avpr1b^Grin1-/-^ (B-F). Areas of Avpr1b gene expression are located between the solid arrows in A and B. C-D. Magnifications of FC (C) and CA2 (D) from B. Solid lines indicate boundaries of Avpr1b gene expression. Solid arrows indicate cells with Avpr1b expression and no Grin1. Dotted arrows indicate cells with remaining Grin1 within the region. E-F. Double in situ hybridization of Grin1 (red) and GAD (green). Solid lines indicate boundaries of the FC (E) and CA2 (F). Solid arrows indicate GAD positive cells with Grin1 expression within these regions. Dotted arrows indicated GAD negative cells with Grin1 expression. G.) Quantification of Grin1 intensity in WT, Avpr1b^Grin1+/-^, Avpr1b^Grin1-/-^. Avpr1b^Grin1-/-^ mice have a significant reduction in Grin1 within the FC and CA2. * indicate p< 0.01 compared to WT and Avpr1b^Grin1+/-.^ Data presented as individual data pints overlaid on means and SD.

#### Normal growth and development in Avpr1b^Grin1-/-^ animals

The transgenic cross did not result any abnormal breeding occurrences and the ratio of the three genotypes was as expected given our breeding strategy [expected: wildtype: 50%, heterozygotes: 25%, knockouts 25%]. Specifically, Avpr1b^Grin1-/-^ males were 8%, Avpr1b^Grin1-/-^ female were 10%, Avpr1b^Grin1+/-^ males were 13%, Avpr1b^Grin1+/-^ females were 16%, WT males were 28% and WT females were 25%, compared to expected (χ^2^,df = 1.15,f. p= 0.5). Litters were normally sized (6.9±2.3) and did not significantly differ in size compared to the Avpr1b-Cre line (Kolmogorov-Smirnov=0.04, p= 0.99).

### Behavioral Results from Avpr1b^Grin1-/-^

#### Resident Intruder Aggression

Experimental mice were tested using the resident-intruder paradigm once a day on three non-consecutive days (Figure 6A). There was no difference in the likelihood to attack an intruder (X^2^=4.4, df=2, p=0.1, Figure 6B.). A two-way repeated measures ANOVA indicated no difference in the duration of aggressive behaviors across trials [F(1.3, 30.6 =0.83, p=0.4) or across genotypes [F(2, 23)=0.5, p=0.9; Figure 6C-D]. Similarly, two-way repeated measures ANOVA indicated no difference in the frequency of aggressive behaviors across trials [F(1.3, 31.5)=0.7, p=0.4] or across genotypes [F(2, 23)= 0.04, p=0.9]. Non-aggressive social interaction toward the intruder was not significantly different between the groups, although a trend for reduced interaction was observed [F(2,23)= 3.3, p=0.057]. All groups showed a habituation to sniffing the intruder over multiple testing sessions (Figure 6G-H). Two-way repeated Measures ANOVA (Trial × Genotype) revealed a significant effect of Trial [F(1,42)=7.49, p=0.002]. Post-hoc Tukey’s multiple comparisons revealed Trial 1 sniffing duration was significantly higher than all other trials [compared with Trial 2 (p=0.001), Trial 3 (p=0.04)].

**Figure 6.**
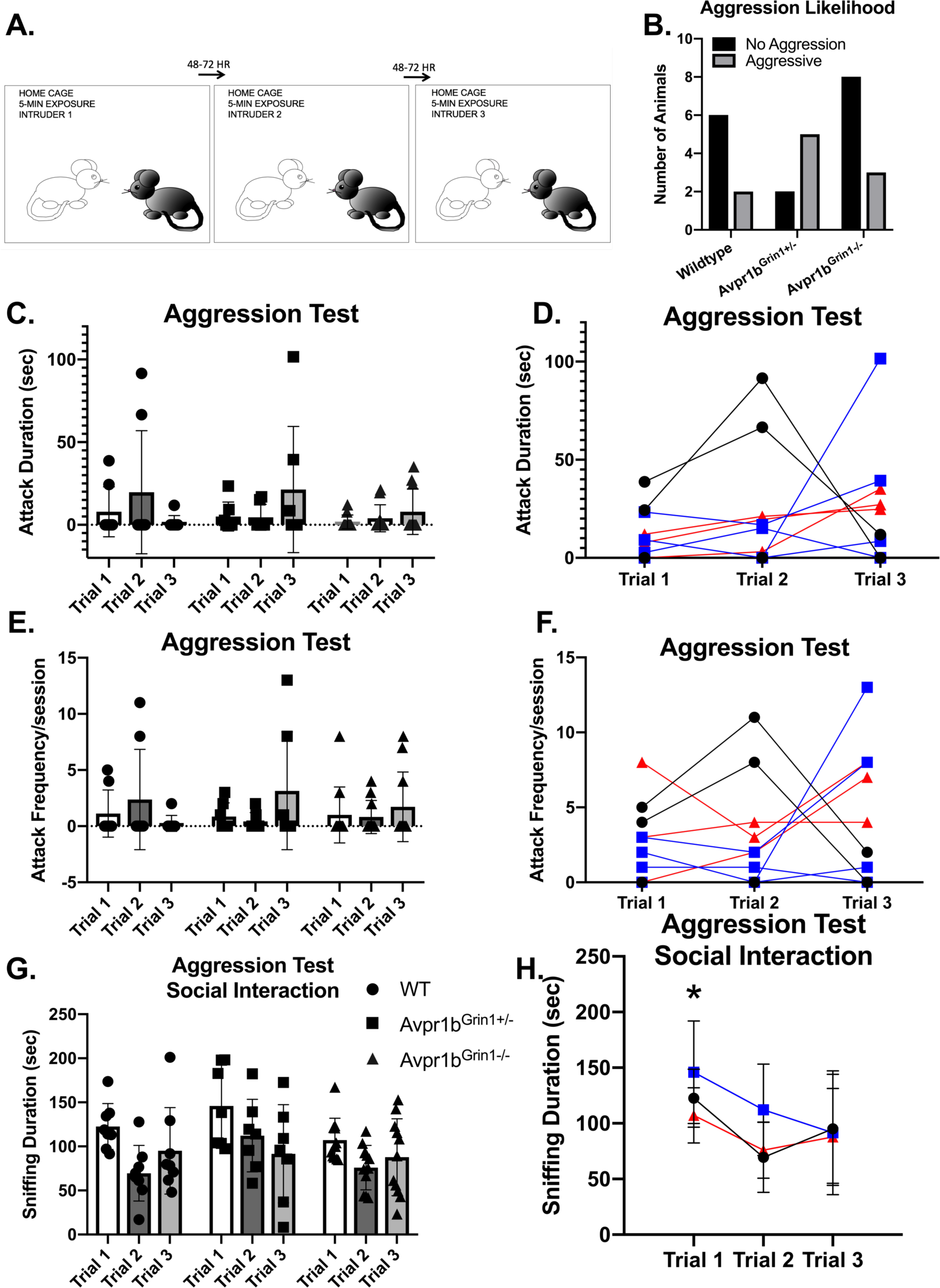
Aggressive Behavior in Avpr1b^Grin1-/-^ Mice. A. Depiction of experimental design. Each mouse was tested on 3 occasion separated by 48-72 hours, each time with a novel intruder. B. Likelihood to attack. The number of mice that exhibited any aggressive behavior across the 3 sessions (black bars) and those that never exhibited any aggressive behaviors (gray bars). No significant difference in ratio was observed. C-F. Aggression Behaviors. Aggression duration (C) and Frequency (E) data presented as individual data points overlaid on means ± SD. D,F) Individual animals tracked across sessions. Wildtypes are represented as black circles and lines. Avpr1b^Grin1+/-^ are represented as blue boxes and lines. Avpr1b^Grin1-/-^ are represented as red triangles and lines. G-H. Sniffing behavior during the aggression test. G.) All data presented as individual data points overlaid on means ± SD. H. Grouped data to illustrate effect of Trial. Two-way ANOVA (Trial X Genotype) revealed a main effect of Trial [F(1,42)=7.49, p=0.002]. Post-hoc Tukey’s multiple comparisons revealed Trial 1 sniffing duration was significantly higher than all other trials. * p< 0.05 Trial 1 vs Trial 2 and 3.

#### Social Memory Testing

##### Social Recognition: Novel-Familiar

No differences were observed in social recognition testing (Fig 7A-B). Two-way repeated measures ANOVA (Genotype × Trial) revealed a main effect of Trial [F (1,24) =75.7, p< 0.0001], with no effect of genotype (p=0.9) on sniffing duration in the social recognition novel-familiar test. Avpr1b^Grin1-/-^ showed similar amounts of sniffing duration in the initial investigation of the stimulus mouse (Sidak’s multiple comparison, WT vs Avpr1b^Grin1-/-^, p=0.9, Avpr1b^Grin1-/-^ vs Avpr1b^Grin1+/-^, p=0.9). A ratio score was calculated to control for individual variability in initial investigation in Trial 1 (Trial 2 -Trial 1/Trial 1). A one-way ANOVA revealed no significant difference between the genotypes [F (2,24)= 0.4, p= 0.6]. Sniffing frequency was unaffected by genotype [F (2,24)= 1.4, p= 0.25] or trial [F (1,24)= 3.1, p= 0.09].

**Figure 7.**
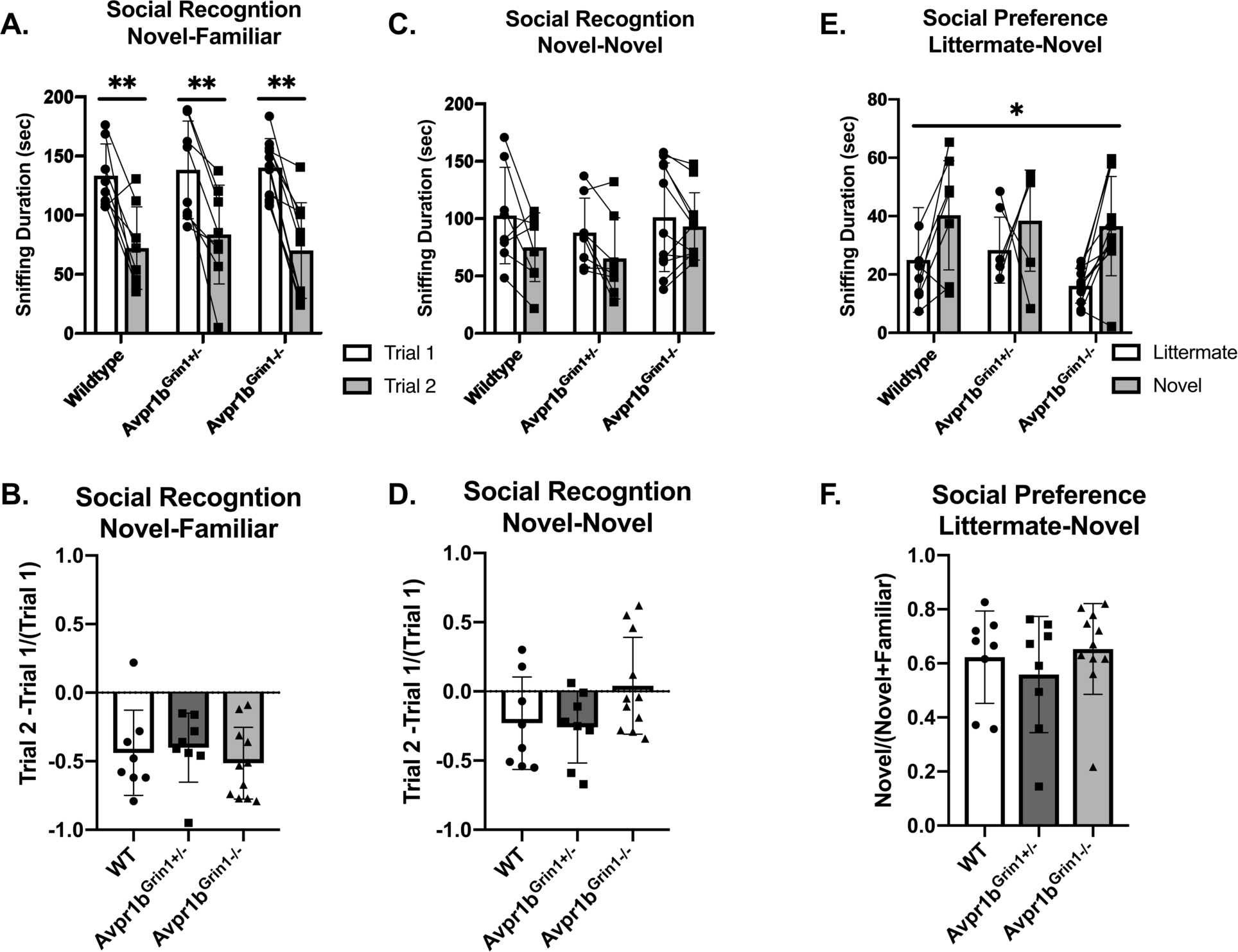
Social Memory is Unaffected by Removal of NMDA Receptor from Avpr1b Neurons. A. Short term (30 min) social recognition measured by repeatedly exposing the subject to the same stimulus animal across two trials (Novel-Familiar). 2-Way ANOVA showed a main effect of trial [F (1,24) =75.7, p< 0.0001]. Post-hoc tests show that within each genotype there is a significant difference between the two trial. ** p < 0.01 B. Ratio of Change between Trial 1 and Trial 2. One-way ANOVA revealed no differences. C. Non-specific recognition response is measured by exposing the subject to different stimulus animals across the two trials (Novel-Novel). 2-Way ANOVA showed a main effect of Trial [F(1,24)=10.7, p=0.003). Posthoc comparisons confirm that no group had a significant difference between trials. D. Ratio of Change between Trial 1 and Trial 2. One-way ANOVA revealed no differences. E. Social novelty preference compares interaction with a littermate and an age- and sex-matched novel animal when presented simultaneously in a 3-chamberd cage. Repeated measures ANOVA revealed a significant main effect of Stimulus Mouse [F(1,20)= 11.1, p=0.003], but no effect of genotype. F. Ratio of Novel sniffing to Total Sniffing. One-way ANOVA revealed no differences. All data are presented as individual responses, with means ± SD overlaid.

##### Social Recognition: Novel-Novel

To determine whether the reduction in Trial 2 was specific to the stimulus mouse, the same protocol was repeated but a novel mouse was presented in the second trial (Figure 7 C-D). We see no difference between the groups [F(2,24)= 0.8, p =0.4], however there was a main effect of Trial [F(1,24)=10.7, p=0.003). Post-hoc comparisons confirm that no group had a significant difference between trials. A one-way ANOVA of the ratio scores revealed no significant differences between the genotypes in performance [F(2,24= 2.6, p=0.09]. Furthermore, there were no differences observed in the initial investigation in Trial 1 [F (2,24)= 0.3, p=0.72].

#### Social Novelty Preference

The social novelty preference design measures the subject’s choice to investigate a novel animal more in the presence of a very familiar animal, in this case a male sibling with which they have continuously been housed. The stimulus mouse was an age-, weight- and sex-matched group housed animal from the same strain (Figure 7E-F). For the sniffing behavior, a repeated measures ANOVA (Genotype × Novelty), revealed a main effect of Novelty [F(1,24)= 12.2, p =0.002]. A ratio score for the investigation time was calculated to determine preference for the novel animal over the littermate (Novel Animal/(Novel Animal + Littermate). There was no significant difference between the genotype in the ratio score (Kruskal-Wallis = 1.03, p= 0.59). Furthermore, there was no difference in the total amount of investigation (Kruskal-Wallis = 2.5, p= 0.28). The total time spent in the side of the chamber yielded similar results. There was a main effect of novelty [F(1,24)= 6.4, p =0.01], with no effect of genotype [F(2,24)=0.7, p= 0.5). The ratio score did not differ based on genotype (Kruskal-Wallis= 0.35, p = 0.8).

#### Social Habituation-Dishabituation

The novel animal trial of social habituation-dishabituation test measures the ability of the subject to recognize an individual after a short exposure and short interval as well as discriminate between individual stimulus animals presented independently. All groups performed similarly in the social habituation-dishabituation test (Figure 8A.). This task presented a new kind of stimulus animal (intact males from the same species). Two-way repeated measures ANOVA (Trial × Genotype) revealed a significant effect of Trial [F(3.0,72.2)=21, p<0.001], but not genotype [F(2,24)=1.1, p=0.4]. Post-hoc Tukey’s multiple comparisons revealed Trial 1 sniffing duration was significantly higher than all other trials [compared with Trial 2 (p=0.005), Trial 3 (p<0.001), Trial 4(p<0.001), Novel (p=0.04)]. Trial 2 was also significantly higher than Trial 4 (p=0.003). A planned post-hoc comparison of Trial 1 confirmed no difference in the initial investigation of the new stimulus mouse type. (WT vs Avpr1b^Grin1+/-^, p = 0.39, WT vs Avpr1b^Grin1-/-^ p = 0.52). Together these indicate a short-term behavioral habituation to the stimulus animal. A robust dishabituation response was observed upon the presentation of the novel stimulus mouse revealed by comparing Trial 4 and the Novel Trial (p<0.001).

**Figure 8.**
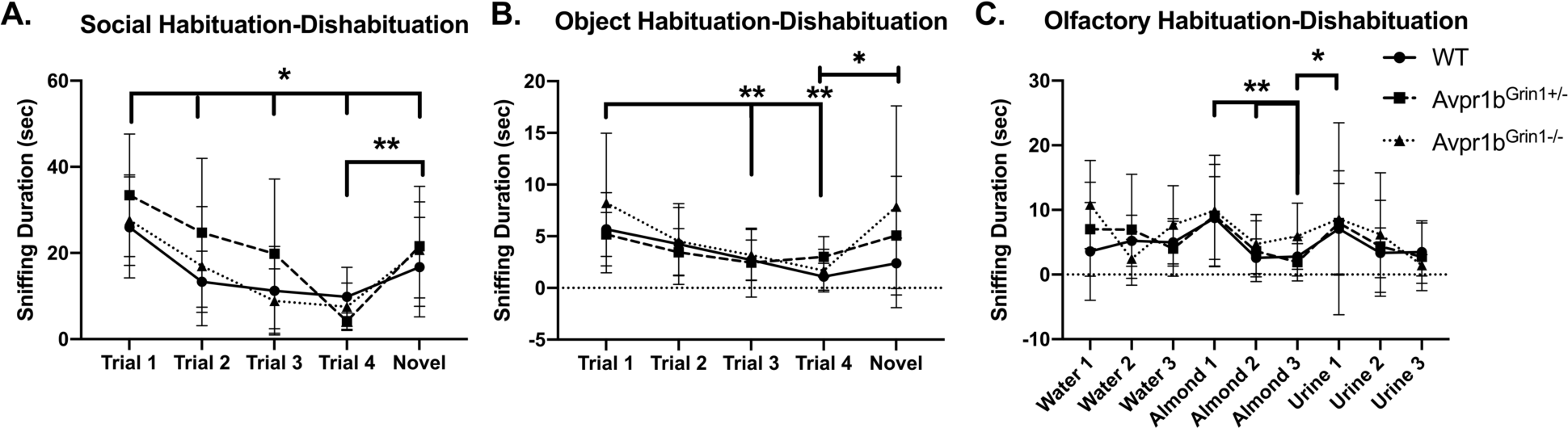
Habituation to Social and Non-social Cues is Unaffected by Removal of NMDA Receptor from Avpr1b Neurons. A. Social Habituation-Dishabituation. Two-way repeated measures ANOVA (Trial × Genotype) revealed a significant effect of Trial [F(4,96)=20.89, p<0.001]. Post-hoc Tukey’s multiple comparisons revealed Trial 1 sniffing duration was significantly higher than all other trials *p< 0.05. A robust dishabituation response was observed upon the presentation of the novel stimulus mouse revealed by comparing Trial 4 and the Novel Trial, ** p<0.001. B. Object Habituation-Dishabituation. Two-way repeated measures ANOVA (Trial X Genotype) revealed a significant effect of Trial [F(4,96)= 7.2, p<0.001]. Post-hoc Tukey’s multiple comparisons revealed Trial 1 sniffing different from Trial 3 and 4, **p < 0.001. A robust dishabituation response was observed upon the presentation of the novel stimulus mouse revealed by comparing Trial 4 and the Novel Trial, * p<0.002. C. Olfactory Habituation-Dishabituation. Two-way repeated measures ANOVA (Trial X Genotype) revealed a significant effect of Trial [F (8,192)=6.1, p<0.001]. Post-hoc Tukey’s multiple comparisons revealed significant decreases between the initial almond presentation and subsequent 2 trials (** p=0.001). Significant dishabituation was observed between the third almond presentation and the first urine presentation (* p=0.04).

#### Object Habituation-Dishabituation

All groups performed similarly in the object habituation-dishabituation test (Figure 8B). There was no difference in the initial interaction time. Two-way repeated measures ANOVA (Trial X Genotype) revealed a significant effect of Trial [F(2.2,53.1)= 7.3, p<0.001], but no effect of genotype [F(2,24)=0.8, p=0.4]. Post-hoc Tukey’s multiple comparisons revealed Trial 1 sniffing duration was significantly higher than later trials [compared with Trial 3 (p=0.001), Trial 4 (p<0.001)]. Together these indicate a short-term behavioral habituation to the presented object. A robust dishabituation response was observed upon the presentation of the novel object revealed by comparing Trial 4 and the Novel Trial (p=0.0025).

#### Olfactory Discrimination

All groups performed similarly in the olfactory habituation-dishabituation test (Figure 8C). There was no difference in the initial interaction time. Two-way repeated measures ANOVA (Trial X Genotype) revealed a significant effect of Trial [F(8,192)=6.1, p<0.001], but not genotype [F(2,24)=0.4, p= 0.7]. Post-hoc Tukey’s multiple comparisons revealed significant decreases between the initial almond presentation and subsequent trials (Almond Trial 2 p=0.003, Almond Trial 3, p=0.003). Significant dishabituation was observed between the third almond presentation and the first urine presentation (p=0.04).

#### Perseverative and escape behaviors

Since NMDA function in the forebrain has been tied to symptoms observed in animal models of schizophrenia (Uno and Coyle 2019), we analyzed perseverative and escape/avoidance behaviors during the social recognition tests. Perseverative behaviors including digging in the bedding and rearing in the cage showed no significant differences between the groups. Specifically during the social recognition novel-familiar test, two-way repeated ANOVA reported no difference in genotype for digging [F(2,24)=0.5, p=0.58] or rearing [F(2,24)=0.3, p=0.72]. To assess escape/social avoidance behaviors, climbing behavior was defined as climbing up the edge of the behavioral chamber with no access to the presented stimuli (Figure 9C-D). Two-way repeated measures ANOVA revealed that during the social recognition novel-familiar test there was a significant effect of Trial [F(1,24)= 10.23, p=0.004), and a trend for an interaction [F(2,24)=3.0, p=0.06] but no main effect for genotype [F(1,24)= 3.1, p=0.059]. Avpr1b^Grin1-/-^ mice had a significant increase in climbing (Sidak’s multiple comparison adjust p=0.001), while WT (p= 0.3) and Avpr1b^Grin1+/-^ (p= 0.9) did not differ between trials. During the social recognition novel-novel test, there was a significant effect of Trial [F(1,24)= 8.0, p=0.0002], but no longer any effect of genotype [F(2,24)= 2.4, p=0.11] on climbing behavior.

**Figure 9.**
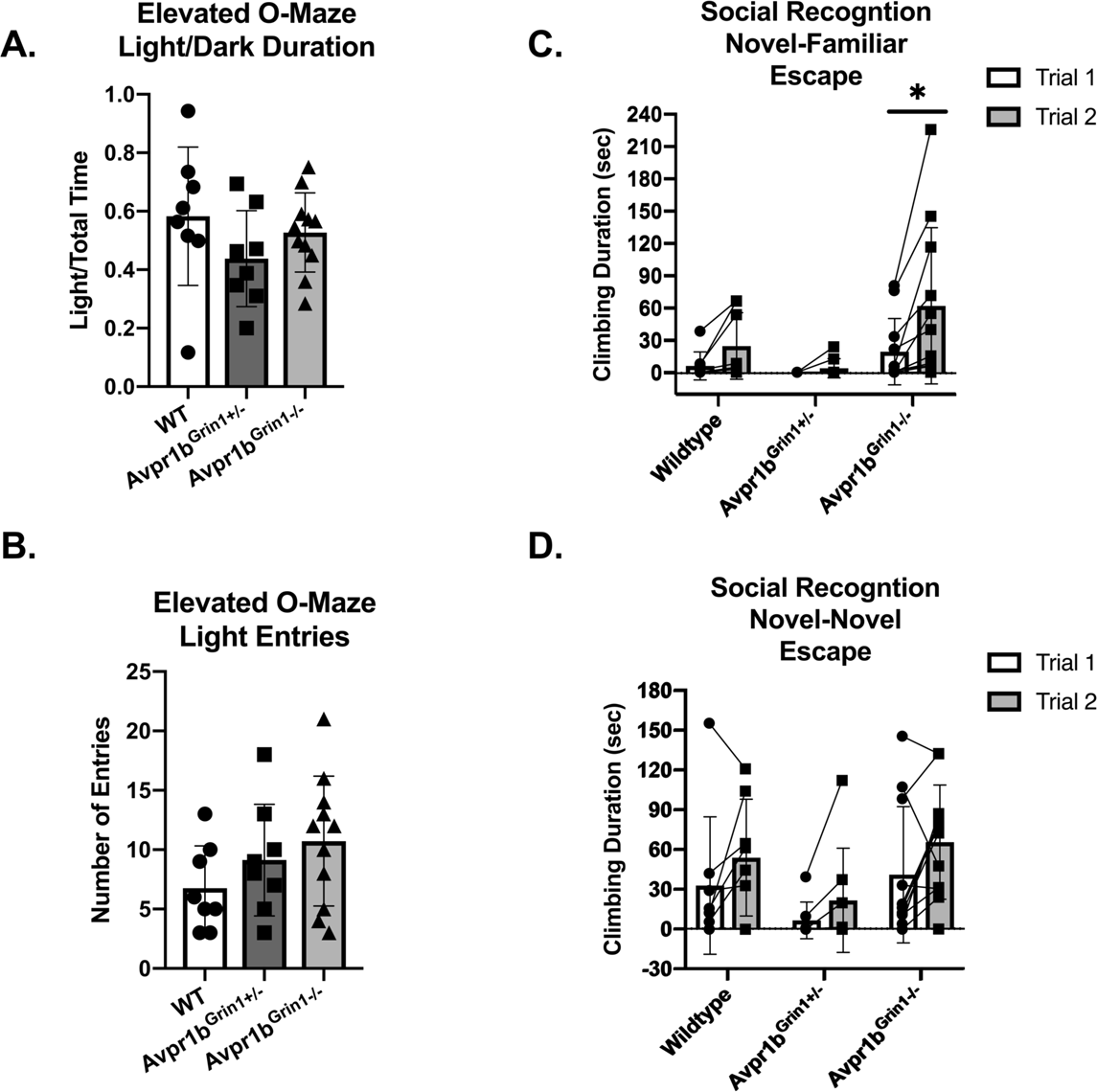
Anxiety-like Behaviors Are Not Disrupted in Avpr1b^Grin1-/-^ Mice. A. Elevated O-Maze. Ratio of duration spent on the light arms. One-way ANOVA indicates no group differences. Locomotor activity indicated by number of entries into either light arm. One-way ANOVA indicates no group differences C-D. Escape behavior (climbing behavior) was measured during the social recognition behavioral tests. C. Two-way repeated measures ANOVA revealed that during the social recognition novel-familiar test there was a significant effect of Trial (F (1,24) = 10.23, p=0.004), but no significance for genotype [F(1,24)=3.1, p=0.059] or interaction [F(2,24)=3.0, p=0.06]. Avpr1b^Grin1-/-^ mice have a significant increase in climbing (*, Sidak’s multiple comparison adjust p=0.001). During the social recognition novel-novel test, there was a significant effect of Trial [F (1,24)=8.0, p=0.0002], but no longer any effect of genotype [F (2,24)=2.4, p=0.11]. All data are presented as individual responses, with means ± SD overlaid.

#### Elevated O-Maze

We measured anxiety-like behavior using the elevated O-maze (Figure 9A-B). There were no significant differences in the number of entries [F(2,24)= 1.6, p=0.21], duration spent in the open arms [F (2,24) = 1.3, p=0.27], or duration of stretching into the light arm (Welch’s W=2.8, p= 0.10). Locomotor activity was unaffected as measured by distance traveled during the test [F(2,24)= 0.66, p= 0.53].

## Discussion

We describe a new transgenic mouse line that placed Cre recombinase expression under the control of the Avpr1b gene promoter. By using this approach, we sought to avoid the physiological and behavioral effects that can accompany unknown genomic insertion sites and off-target Cre-expression that are common in many BAC transgenic lines (Heffner, Pratt et al. 2012, Harno, Cottrell et al. 2013). Gross physiological development is normal in this strain with litter number, sex ratio, and adult weight comparable to C57Bl/6J mice in our lab. We compared heterozygous Avpr1b-Cre mice to their WT siblings and found no differences in social behaviors, confirming that the Cre insertion did not cause off-target behavioral effects in this strain. Avpr1b and Cre mRNA localize to CA2 neurons, while there is minimal expression within the dentate gyrus, CA1 or CA3. Using a transgenic constitutive labeling approach (i.e, cross with a Cre-activatable fluorescent marker mouse line), we observed labeled neurons within the dorsal and ventral CA2, olfactory bulbs, caudate-putamen. Although we propose this model will be useful for CA2 neuronal targeting, caution should be taken with constitutive approaches. For example, we do not see Avpr1b expression by in situ hybridization histochemistry in the adult caudate-putamen, presumably because transient expression of Avpr1b driving Cre earlier in development activated the fluorescent marker gene.

Crossing this Avpr1b-Cre line with a transgenic line with a floxed Grin1 gene, resulted in the removal of functional NMDA receptors from Avpr1b neurons. We assessed this line in a battery of behavioral tests measuring memory, social and emotional behaviors. Surprisingly, a lack of NMDA in Avpr1b (including most CA2) neurons showed minimal effect on any of the behaviors assessed in this study. This indicates that NMDA-dependent mechanisms may not be required for Avpr1b neuronal plasticity and Avpr1b-dependent behaviors.

Avpr1b KOs show a robust deficit in aggressive behavior in both males and females (Wersinger, Ginns et al. 2002, Wersinger, Kelliher et al. 2004). Replacement of Avpr1b into the dorsal CA2 of Avpr1b KO males results in the rescue of aggressive behavior (Pagani, Zhao et al. 2015). Furthermore, the acute activity of dorsal CA2 neurons has also recently been shown to be required for typical expression of aggression (Williams Avram, Lee et al. 2016, Leroy, Park et al. 2018). The exact mechanism through which Avpr1b activity may be contributing to the role of CA2 in aggression remains poorly understood. Aggression is often considered an innate behavior but has an important learned component. Aggressive behavior skills are developed by male mice during adolescence, and change based on experience. NMDA receptor activity within the ventral hippocampus, where there are abundant Avpr1b neurons, is critical for typical aggressive behavior expression (Chang, Su et al. 2018). Surprisingly, we do not observe any differences in aggressive behavior in the Avpr1b^Grin1-/-^ mice, either in the likelihood to exhibit the behavior or in the duration or frequency of attacks.

Avpr1b KO mice have deficits in short-term social recognition and social habituation (Wersinger, Ginns et al. 2002, Stevenson and Caldwell 2012, Williams Avram and Cymerblit-Sabba 2017). Similarly, disruption of CA2 neuronal activity chronically and acutely can inhibit short-term social recognition (Hitti and Siegelbaum 2014, Stevenson and Caldwell 2014, Meira, Leroy et al. 2018). Although the activity of pyramidal neurons of the CA2 is required for social recognition, NMDA-dependent molecular plasticity within the pyramidal neurons may not be required. Similar to our findings, Finlay et al. show that genetic removal of NMDRs from the dorso-lateral hippocampus (inclusive of CA2 and CA3 neurons) in adulthood resulted in no effect on social novelty preference, which requires social memory (Finlay, Dunham et al. 2015). Furthermore, although pyramidal CA2 neurons do not respond to standard long-term potentiation inducing protocols, it was recently observed that input-timing dependent plasticity can occur, but that this change in pyramidal neuron response is driven by long-term depression of adjacent parvalbumin (PV) interneurons in the CA2 region (Dudek, Alexander et al. 2016, Leroy, Brann et al. 2017). Blocking delta opioid receptors, which are found on PV interneurons in the CA2, was able to block the depression in the PV interneurons, the potentiation in the pyramidal neurons and short-term social memory (Leroy, Brann et al. 2017). These elegant studies provide a potential mechanism through which CA2 neurons may not require NMDA-dependent plasticity.

With the exception of a subtle effect in temporal order memory (DeVito, Konigsberg et al. 2009), Avpr1b KO mice do not show deficits in non-social memories (Wersinger, Ginns et al. 2002, Wersinger, Kelliher et al. 2004). Similarly, we did not observe any object of olfactory memory deficits in Avpr1b^Grin1-/-^ mice. Although NMDA signaling in other regions of the hippocampus have been shown to be critical for non-social memories. The constitutive CA1 Grin1 KOs show a strong deficit in object memory and olfactory memory. Interestingly, these deficits are rescued by exposure to enriched conditions (Rampon, Tang et al. 2000). Similarly, CA3 Grin1 KOs exhibit deficits in acquisition and recall of associative memories (Nakazawa, Quirk et al. 2002). Notably, inducible removal of Grin1 from CA1 neurons during adulthood results in deficits in the acquisition of spatial memory (Shimizu, Tang et al. 2000), but no effect on recall. This deficit is only observed in inducible models as constitutive removal of Grin1 resulted in no alteration in memory performance (Mei, Li et al. 2011). These data suggest a complex context, and memory-type specific role for NMDA receptors within hippocampal pyramidal neurons.

Reductions in NMDA signaling either pharmacologically or genetically have generally been found to disrupt social interactions. NMDA antagonists such as PCP can reduce social interactions and social recognition performance (Neill, Barnes et al. 2010, Zimnisky, Chang et al. 2013). Furthermore, NMDA hypomorphs have reduced social interaction (Mohn, Gainetdinov et al. 1999, Duncan, Moy et al. 2004, Duncan, Inada et al. 2009, Halene, Ehrlichman et al. 2009, Mielnik, Horsfall et al. 2014). Genetic removal of NMDA receptors throughout the forebrain during adulthood results in impairments in social approach, as well as reduced long-term social recognition (Jacobs and Tsien 2017). Similarly, removal of Grin1 from dorso-lateral hippocampus in adulthood results in decreases in social approach (Finlay, Dunham et al. 2015). However, we observed no effect on social interaction in any of our tests. Our data are in agreement with previous studies of Avpr1b KO mice, which do not show reductions in social interaction or social approach behavior (Wersinger, Ginns et al. 2002, Yang, Scattoni et al. 2007). Furthermore, no studies investigating the chronic or acute disruption of CA2 neurons have reported any changes in social interactions or social approach (Hitti and Siegelbaum 2014, Stevenson and Caldwell 2014, Piskorowski, Nasrallah et al. 2016, Smith, Williams Avram et al. 2016, Leroy, Brann et al. 2017, Meira, Leroy et al. 2018).

## Conclusions

To our knowledge, this is the first report to indicate that NMDA function is not required in neurons known to play a critical role in memory performance. Given the lack of behavioral effects, it is tempting to conclude that NMDA receptors on Avpr1b neurons in the hippocampus have no role in social behaviors or social memory. However, social approach and social recognition performance can be disrupted by manipulation of several distributed brain areas including olfactory processing regions, the paraventricular nucleus, and the amygdala; some of which contain low numbers of Avpr1b neurons. Furthermore, it is possible that the constitutive removal of NMDA receptors results in compensatory changes in these, or other, neurons to allow for performance of these important behavioral functions. For example, it is known that the Avpr1b-mediated enhanced EPSC response observed in dorsal CA2 neurons can be blocked by NMDA receptors blockade, but also blocked by interrupting other intracellular calcium signaling mechanisms. Avpr1b-expressing neurons may have adapted the function of these mechanisms to permit the typical response. CA2 neurons have distinct calcium buffering properties and are uniquely encased with a perineuronal net compared to neighboring CA1 and CA3 neurons. Further investigation using our novel Avpr1b-Cre transgenic mouse line will help elucidate mechanisms within Avpr1b neurons and how they contribute to social aggression and memory

## Acknowledgments

This research was supported by the intramural research program of the NIMH (ZIAMH002498). We would like to thank the members of the Section for Neural Gene Expression that helped over the years to collect this data. The NIMH Transgenic Core for its services with transgenic animals. The Systems Neuroscience Imaging Resource allowed us to use imaging equipment and software.

